# Aglycone polyether ionophores as broad-spectrum agents inhibit multiple enveloped viruses including SARS-CoV-2 in vitro and successfully cure JEV infected mice

**DOI:** 10.1101/2020.10.27.354563

**Authors:** Jia-Qi Li, Minjian Huang, Ya-Nan Zhang, Ran Liu, Zhe-Rui Zhang, Qiu-Yan Zhang, Yong Wang, Jing Liu, Zixin Deng, Bo Zhang, Han-Qing Ye, Tiangang Liu

## Abstract

Infections with zoonotic viruses, such as flaviviruses, influenza virus, and the SARS-CoV-2 pandemic coronavirus constitute an increasing global risk. Hence, an urgent need exists for the development of broad-spectrum antivirals to prevent such outbreaks. Here, we show that the maduramycin and CP-80,219 aglycone polyether ionophores exhibit effective broad-spectrum antiviral activity, against various viruses, including Japanese encephalitis virus (JEV), Dengue virus (DENV), Zika virus (ZIKV), and Chikungunya virus (CHIKV), while also exhibiting promising activity against PR8 influenza virus and SARS-CoV-2. Moreover, liposome-encapsulated maduramycin and CP-80,219 provide full protection for mice from infection with JEV *in vivo*. Mechanistic studies suggest that aglycone polyether ionophores primarily inhibit the viral replication step without blocking endosome acidification to promote the fusion between viral and cellular membranes. The successful application of liposomes containing aglycone polyether ionophores in JEV-infected mice offers hope to the development of broad-spectrum antiviral drugs like penicillin back to 1940s.

## Introduction

Recently, zoonotic viruses have caused various infectious disease outbreaks across the world, posing a serious threat to human and animal health, For example, the current severe acute respiratory syndrome coronavirus 2 (SARS-CoV-2) pandemic has caused over 40 million infections and more than 1.14 million deaths^1^. In fact, approximately 89 % of the 180 recognized RNA viruses with the potential to harm humans are zoonotic^2^, and most of them are enveloped to facilitate adaptation to different hosts. Flaviviruses as a major category of zoonotic viruses comprise more than 70 members, including Japanese encephalitis virus (JEV), Dengue virus (DENV), West Nile virus (WNV), and the most recent explosive epidemic of Zika virus (ZIKV) in the Americas. These viruses are primarily transmitted by arthropod vectors, such as mosquitoes, midges, and sand flies and infect millions of people annually, resulting in thousands of deaths^3, 4^. Flavivirus Infections present with varying symptoms, ranging from flu-like illnesses and fevers to potentially lethal haemorrhagic fevers, and neurological complications, include encephalitis, microcephaly and Guillain–Barré syndrome^5, 6^.

Several antivirals and vaccines targeting flaviviruses, including the Hepatitis C (HCV) nucleoside inhibitor sofosbuvir^7^, as well as vaccines for Yellow fever virus (YFV) and JEV^8^, have been licensed and are commercially available in various countries, but the high mutation rate associated with RNA viruses makes development of effective options challenging. Specifically, RNA viruses have an inherent error-prone process that facilitates the accumulation of mutations during genome replication. This high mutation capacity is what makes the generation of a universal vaccine or drug for RNA viruses, a seemingly impossible task, and it is unlikely to be adequate for ‘one bug–one drug’ approach to antiviral drug development. Moreover, the strategies used for the development of vaccines and antivirals are not amenable to emerging viruses or to the mutations of existing viruses. Hence, the development with broad-spectrum antivirals is a complex, yet warranted task to address the emerging and reemerging virus outbreaks.

Although effective treatment options for RNA viruses remain largely elusive, greater strides have been made in the fight against bacteria, fungi, and parasites, in the form of broad-spectrum antibiotics, such as like penicillin, candicidin and avermectin. Meanwhile, the long-term focused research of our team, in the veterinary and poultry antibiotics industry, revealed that microbial whole fermentation broth, which is used as the primary component of antibiotics administered to animals and poultry, contributes to the reduced occurrence of viral infection, compared with the use of pure antibiotics. Hence, these fermentation broths might contain previously neglected compounds capable of eliciting antiviral activity. Therefore, in the current study, we obtained supernatants from 13 kinds of actinomycetes, originating from current industrial strains used in antibiotic production, and screened their antiviral activity using the JEV platform. Results show that aglycone polyether ionophores exhibit effective antiviral activity, with the half-maximum effective concentration (EC_50_) values of maduramycin and CP-80,219 of 11 nM and 6 nM, respectively. *In vivo* studies demonstrated that liposome encapsulation of maduramycin and CP-80,219 provided full protection of mice from JEV infection. Moreover, mechanistically, maduramycin primarily hinders the critical steps of virus replication cycle, not interfere with viral attachment and entry and not affect endosome acidification. Furthermore, we found that maduramycin and CP-80,219 also exhibit significant antiviral activity against ZIKV, DENV, CHIKV, and PR8 influenza virus, and especially promising active against the SARS-CoV-2 pandemic coronavirus, with EC_50_ of 77.7 nM and 8.7 nM, respectively. These data suggest that maduramycin and CP-80,219 aglycone polyether ionophores have the potential to prevent an organism from becoming a possible transmission or storage host, and block the transmission between humans and animals.

## Results

### Screening of fermentation products of actinomycetes for anti-JEV virus activity

First, we fermented 13 industrial actinomycete strains collected from our lab, including two strains producing polyethers, five strains producing macrolides, and three strains producing aminoglycosides, one strain producing cyclic lipopeptide, one strain producing ε-poly-lysine and 1 strain producing tetracycline (Fig. 1a, more detailed information see Supplementary Table 1). In addition to these primary products, these strains are also known to produce a variety of other natural compounds. These actinomycete strains were cultured by three different fermentation mediums to obtain 39 supernatant samples according to strict microbial fermentation environmental conditions, as showed in Fig 1a. We then tested the 39 supernatant samples for their antiviral activity on the JEV platform (Fig. 1b-d), we showed that the supernatants of J1-005 (producing ε-poly-lysine) fermented in three culture mediums and the supernatant of J1-002 fermented in A medium were high cytotoxicity for Vero cells and not detected their antiviral activity, and other samples in medium A and C are also not able to inhibit JEV virus replication. Surprisingly, the supernatants of two polyether-producing actinomycetes, fermented products of which primarliy included maduramycin and salinomycin in medium B, reduced virus production with titer approximately 3 log10 and 1 log10 compared to blank treatment in a plaque reduction assay. These results displayed that polyether ionophores could show promising antiviral activity.

**Fig. 1.**
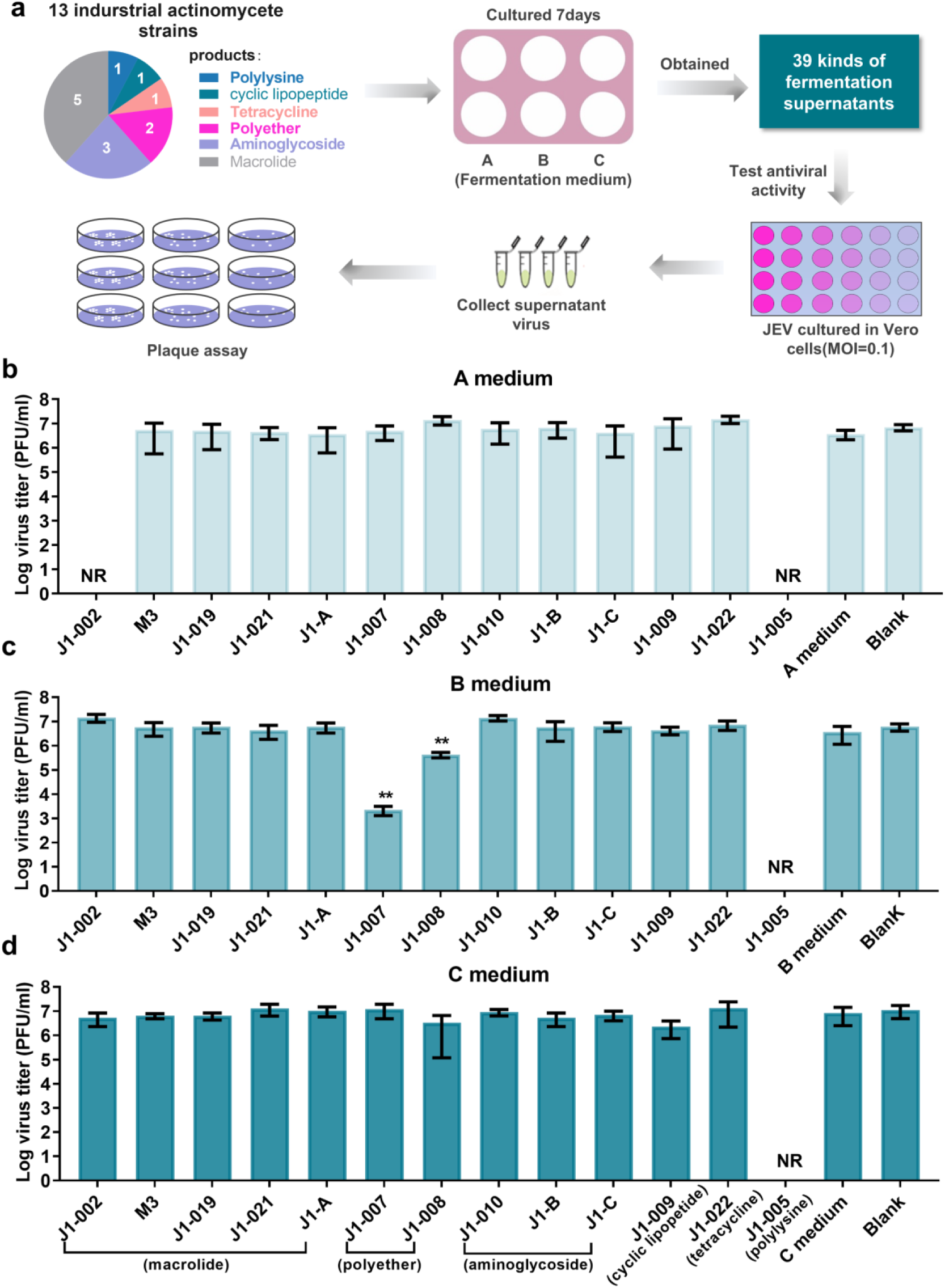
The screening anti-JEV activity for supernatants of 13 actinomycetes strains. a) The workflow of protocol for screening anti-JEV virus activity. b-d) The antiviral activity of 39 kinds of actinomycetes fermentation supernatants on JEV platform. Vero cells infected with JEV (MOI=0.1) in the presence of 10 µL of each sample for 24 h, then evaluated activity by a plaque reduction assay (n=3). NR represents negative results with high cytotoxicity for Vero cells. **, P < 0.01. MOI: multiple of infection.

### Six polyether ionophores exhibit highly active against JEV

Considering that the supernatants of two polyether-producing actinomycetes exhibited promising activity, and the antiviral capability of maduramycin-producing industrial strain supernatant was greater than that of salinomycin-producing strain, combinied with our previous knowledge regarding the biological activity of aglycone polyether ionophores (including maduramycin)^9^, we postulated that aglycone polyether ionophores might exert parallel antiviral activity. Therefore, we first prepared and purified nonaglycone polyether salinomycin as well as five aglycone polyether products, including maduramycin, A-130-A, nanchangmycin, endusamycin and CP-80,219 from their produced actinomycetes, respectively (Fig. 2a). We then assessed the antiviral activity and toxicity of these polyether ionophores against JEV in Vero cells exposed to a drug dose range for 24 h. Results show that all polyether ionophores were effective against JEV, and a cytotoxicity greater than 40 μM (Fig. 2b-2g). Moreover, CP-80,219 and maduramycin were highly efficacious with an average EC_50_ of 5.6 nM and 10.7 nM, respectively. These data indicate that aglycone polyether ionophores exhibit potent antiviral activity against JEV.

**Fig. 2.**
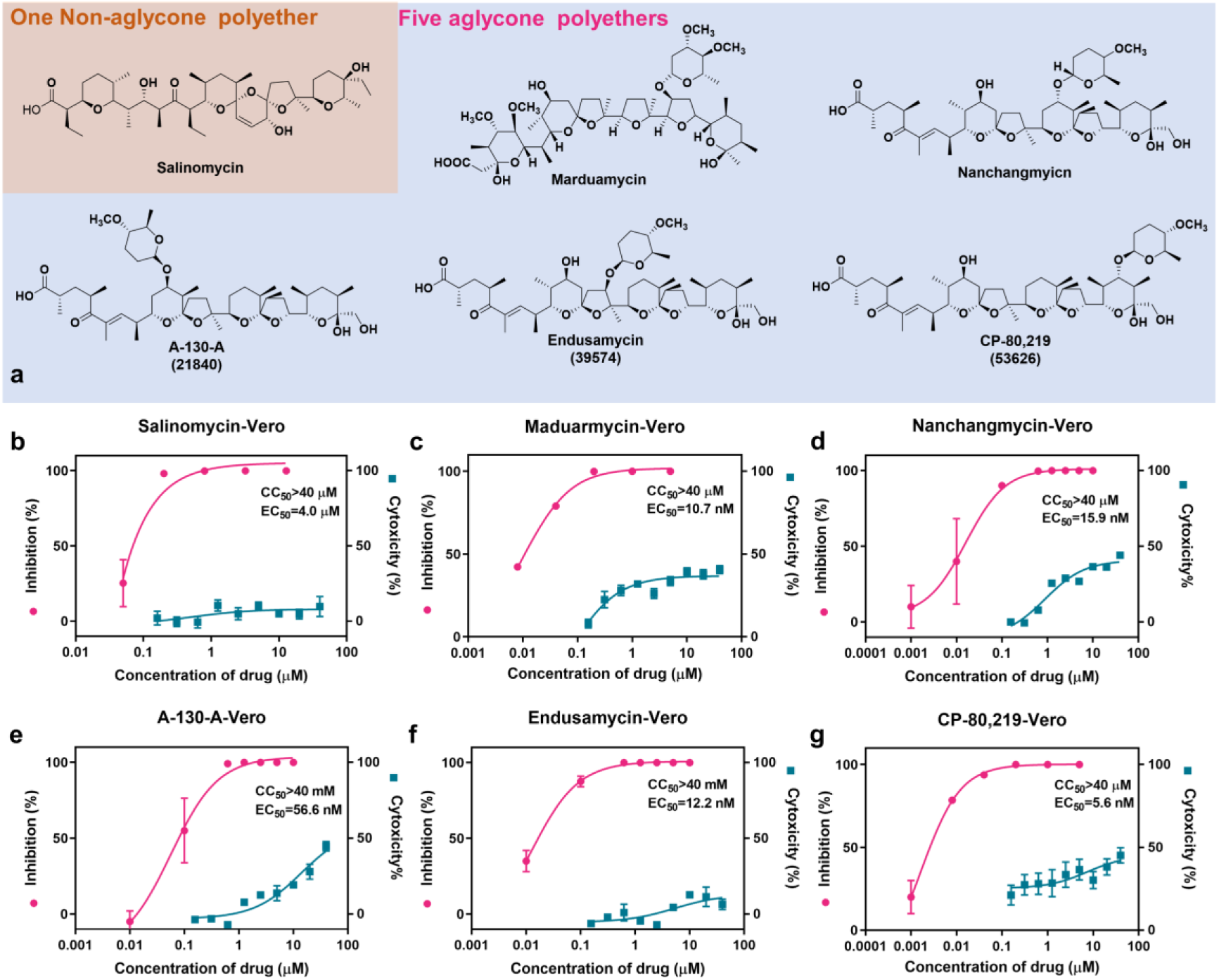
Antiviral activity of six ployether ionophores (salinomycin, maduramycin, nanchangmycin, A-130-A, endusamycin and CP-80,219 against JEV in Vero cells. a) The chemical structures of six polyether ionophores, they are isolated from their actinomycete strains and confirmed by NMR and HRMS. b-g) Vero cells infected with JEV (MOI=0.1, n=3) in the presence of a range of drug for 24 h, after virus replication was measured by plaque assay, and cytotoxicity (n=4) was evaluated by CCK8 assay kit.

### Comparison of antiviral activity between free and liposome-encapsulated maduramycin and CP-80,219 aglycone polyether ionophores

Although maduramycin and CP-80,219 displayed promising biological activity against JEV *in vitro*, studies have shown that the dosage of polyether ionophores in animals is strictly limited^10, 11^. For instance, regarding maduramycin, its maximum dosage for administration to broiler chickens and turkeys is set to 5 mg/kg in feed^12^. Furthermore, following the evaluation of other biological activities of polyether ionophores *in vivo*, toxic effects, including cardiac toxicity and hepatotoxicity, were observed^9, 13^. Meanwhile, encapsulation of drugs in liposomes is known to enhance the therapeutic indices of various agents, primarily through alterations in their pharmacokinetics and pharmacodynamics^14^, such as in the case of the liposome-based paclitaxel formulation (LEP-ETU) for cancers^15^ and doxorubicin encapsulated in liposomes (Doxil) for ovarian and breast cancer^16^. We, therefore, used maduramycin and CP-80,219 with distearoylphosphatidylcholine (DSPC) and cholesterol to form maduramycin (maduramycin-lipid) and CP-80,219 liposome (CP-80,219-lipid) with an average encapsulation rate of 85% and 89%, respectively (Supplementary Fig. 2), and a mean particle size of approximately 100 nm (Fig 3a). Then we investigated whether these liposomes exhibit the same antiviral activity, while reducing cytotoxicity *in vitro* using Vero and BHK cells (Fig. 3b and Fig. 3c). Our results show that the liposome formulations of maduramycin and CP-80,219 maintained the same biological activity in the two host cells with EC_50_ values of up to 3-22 nM and cytotoxicity greater than 40 μM. these results suggest that liposome encapsulation maintains the relatively high antiviral activity of each compound, contributing to the investigation of their biological activity *in vivo*.

**Fig. 3.**
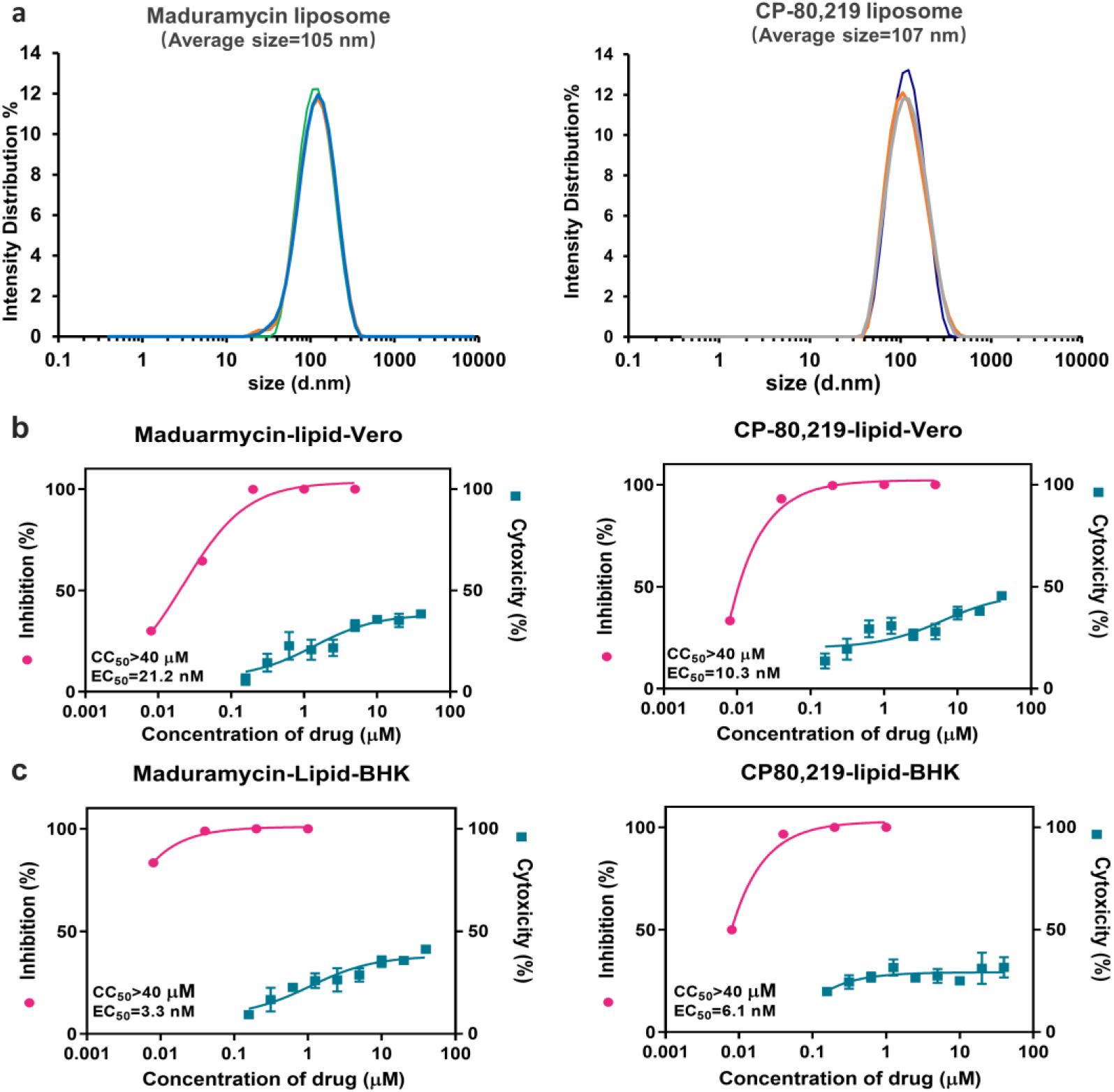
The antiviral activity and cytotoxicity for free and liposome-encapsulated maduramycin and CP-80,219. a) Liposome average size of maduramycin and CP-80,219 was detected by Zetasizer Nano ZS90, with 105 nm and 107 nm respectively. b and c) BHK cells and Vero cells infected with JEV (MOI=0.1, n=3) in the presence of a range of drug for 24 h, after virus replication was measured by plaque assay, and cytotoxicity (n=4) was evaluated by CCK8 assay kit.

### Efficacy of liposome-encapsulated maduramycin and CP-80,219 in mice

Given the efficient *in vitro* antiviral activity of maduramycin-lipid and CP-80,219-lipid against JEV, we next aimed to evaluate efficacy of these liposomes in preventing JEV infection in C57BL/6 mice. To this end we intraperitoneal administrated vehicle (WT group, PBS), 1 mg/kg maduramycin-lipid, or 1 mg/kg CP-80,219-lipid after intraperitoneal infection with 1×10^7^ PFU of mouse-adapted JEV. Subsequently vehicle or drug was administered every 24 h until the end of the study. C57BL/6 mice treated with only maduramycin-lipid or CP-80,219-lipid exhibited slight weight loss through the study. However, this effect was rapidly diminished following vehicle treatment at post-stage, while mortality was observed in JEV-infected mice (Vehicle treatment) at 6 dpi, reaching 100% at 9 dpi. Surprisingly, all ten animals treated with maduramycin-lipid and CP-80,219-lipid formulations survived their exposure to the end of in-life phase. Furthermore, the plasma viral load in all maduramycin-lipid and CP-80,219-lipid-treated groups compared with vehicle-treated groups was found to be significantly decreased with geometric means reduced by ≥ 1.0 log10 (P<0.01) compared to the vehicle-treated group at 1 dpi. Altogether, these results further demonstrate the antiviral activity of maduramycin-lipid and CP-80,219-lipid formulations in an *in vivo* mouse model of JEV infection.

### Aglycone polyether ionophores primarily inhibit the replication stage of JEV life cycle

Studies have shown that polyether ionophores can inhibit virus replication by interfering with the function of viruses and host cells. For example, monensin selectively blocks the secondary glycosylation step of HIV gp160 with the reduction of syncytium formation^17^ and easily induces the alkalization of intracellular organelles to inhibit the fusion between viral and cellular membranes of the late step of HCV entry^18^. Meanwhile, salinomycin prevents the endosomal acidification of host cells and viral matrix protein, thus inhibiting influenza virus infection^19^. To elucidate the mechanism underlying the antiviral activity of aglycone polyether ionophores, Vero cells were treated with 10 µM maduramycin for 1 h or 1 µM maduramycin for 24 h, then acridine orange was used to stain the intracellular vesicles to monitor changes in the pH. Consequently, red fluorescence (representing acidic condition) was observed in the cytoplasm of cells exposed to maduramycin (Supplementary Fig. 1), indicating that aglycone polyether ionophore may not cause cytoplasmic alkalization to block the fusion of viral and cellular membranes.

Next, we explored the effects of aglycone polyether ionophores on viral life cycles. First, we investigated the virucidal activity in depth using time of addition of maduramycin in a plaque assay (Fig. 5A and Fig. 5B). Host cells pretreated maduramycin for 1 h did not inhibit virus replication, and virus titer was not reduced at 1 hpi (hour post-infection). Meanwhile, when host cells are pretreated for 1 h, followed by treatement with the drug for 12 hpi (−1-12), we observed that the viral titer of supernatant nearly was undetectable. A similar result was detected following treatment for cells with the drug for 12 hpi (0-12), without pretreatment. During different periods of drug incubation with 11 h (1-12), 10 h (2-12), 8 h (4-12) and 4 h (8-12), maduramycin substantially reduced virus titer reduction greater than 3 log10 values. Even after 10 h of infection and treated with maduramycin for 2 hpi, the supernatant viral titer was decreased rapidly (>1.5 log10). These data suggest that treatment with maduramycin can significantly inhibit JEV release from host cell. Finally, cells were infected JEV at an MOI (multiple of infection) of 1 for 2 h in the presence of maduramycin or DMSO. At 2 hpi, cells were rinsed, and total RNA was isolated from parallel wells, and Q-PCR analysis was performed. We did not detect any significant alteration in total RNA between treated and untreated cells (Fig. 5c). This finding demonstrates that maduramycin not interfere with the attachment or entry cycle of JEV into the cells.

**Fig. 4.**
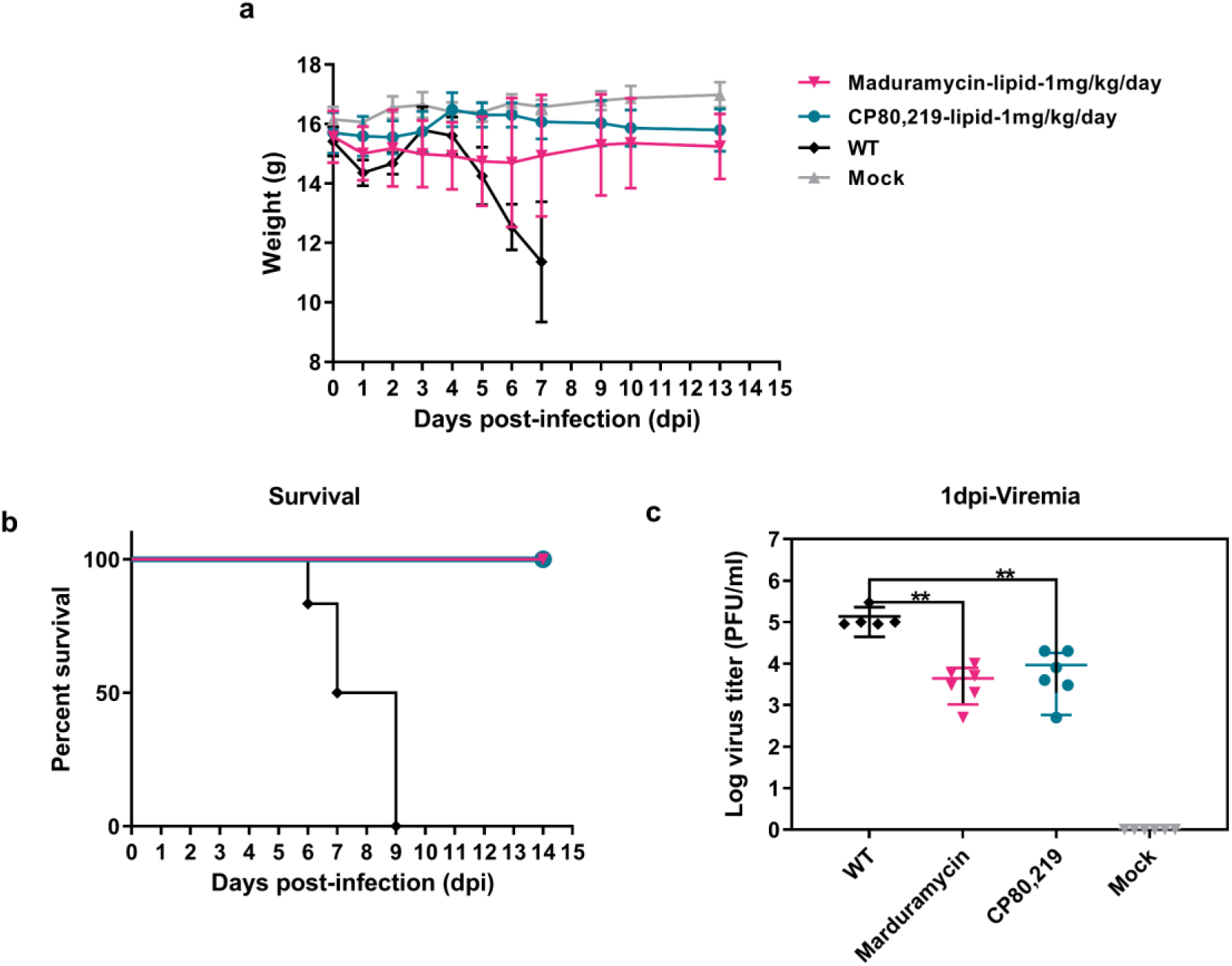
Maduramycin lipsome and CP-80,219 liposome inhibit JEV virus in mice. a) The weight change after maduramycin liposome (1mg/kg, n=6) and CP-80,219 liposome (1mg/kg, n=6) by intraperitoneal administration daily for 14 days. WT group (n=5) as vehicle was infected JEV mice, and Mock group (n=5) was normal mice. b) survival curves for treatment versus vehicle groups assessed by log-rank analysis using Dunnett–Hsu procedure to adjust for multiple comparisons. c) Viremia was determined from blood collected from the eye socket in plaque assay at 1dpi. **, P < 0.01.

**Fig. 5.**
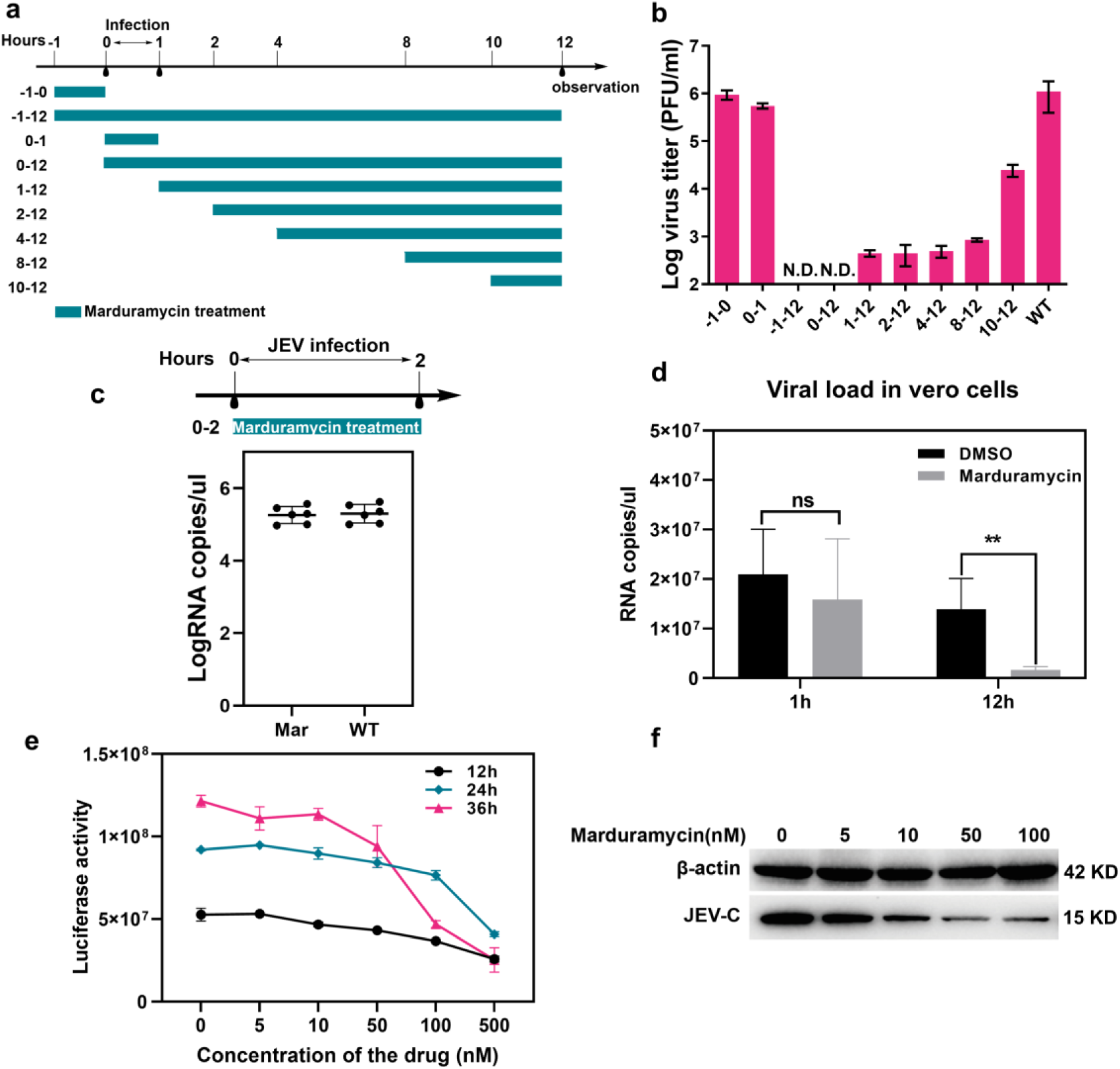
The mechanism of action of polyether maduramycin. a) Time course of virucidal activity: JEV (MOI=10) and Maduramycin (500 nM) were incubated in Vero cells for different time periods in a plaque assay. N.D.: no detected virus in supernatant. b) Time course of virucidal activity were detected by Plaque assay (n=3). c) Vero cells were infected with JEV-WT at a MOI of 1 for 2 h. Infected cells were treated with 0.1% DMSO (solvent control) and Maduramycin (500 nM), At 2 hpi, cells were rinsed and total RNA was isolated from parallel wells, and Q-PCR analysis was detected (n=6). d)Vero cells were infected at a MOI of 1 with wild-type JEV with Maduramycin (500 nM) for 1 h, then the infected cells were washed thrice with PBS, and then 1 mL fresh medium was added to each well, and maduramycin (500 nM) were added for 1 h and 12 h. The RNA copies were detected as described above at 12 hpi (n=4). ns: no significant difference; **, P < 0.01. e) JEV-Rluc-replicon cell line were seeded in 24-well plate with 50000 cells per well, after 16 h, different concentration of Maduramycin was treated, and Rluc signals were detected at different time point. f) Vero cells were infected at an MOI of 0.1 with wild-type JEV for 24 h with different concentration maduramycin (n=3). Cell lysates were analyzed by Western blotting for the presence of the indicated JEV capsid protein (JEV-C). β-actin was used as a loading control. All results from 3 independent experiments.

Subsequently, we analyzed whether maduramycin acts on virus replication process. Cells were infected JEV at a high MOI of 10 and incubated with maduramycin or DMSO for 1 h or 12 h at 37 °C, we then detected JEV genome RNA copies and observed a highly significant effect (P=0.0075) between the maduramycin and DMSO groups (Fig.5D), indicating that maduramycin could affect virus replication. Furthermore, we used JEV-Rluc-replicon cell line and found administration of maduramycin decreased JEV-Rluc-replicon replication in dose- and time-dependent manner (Fig.5e). Consistently, we also observed a dose-dependent reduction in the levels of JEV viral capsid protein when cells were infected JEV with different concentration maduramycin (Fig.5f).

### Maduramycin and CP-80,219 ionophores exert broad-spectrum antiviral activity including the new SARS-CoV-2 coronavirus

Early studies have shown that a variety of polyether ionophores exhibit biological activity against HIV virus^20^. Subsequently, nanchangmycin has been reported to exert broad-spectrum antiviral effects, including against flaviviruses, by inhibiting the invasion of viruses, suggesting that polyether ionophores might have potential antiviral activity as broad-spectrum reagents^21^. To determine whether maduramycin and CP-80,219 also exert broad inhibition of viral replication, we investigated their biological activity against enveloped viruses using a plaque assay (Fig. 6). Accordingly, both the maduramycin and CP-80,219 ionophores were shown to efficiently inhibit the replication of these virus pathogens, including ZIKV, DENV and Chikungunya (CHIKV) (Fig 6a), with maduramycin exhibiting a nanomolar level activity against DENV and CHIKV. Both of aglycone polyether ionophores targeting against PR8 influenza virus exhibited promising inhibitory and low cytotoxicity (Fig. 6b). In addition, maduramycin and CP-80,219 ionophores were effective against the epidemic SARS-CoV-2, with EC_50_ of 70 nM and 9 nM, respectively (Fig. 6c), which are more efficient than the currently reported remdesivir (EC_50_=0.77 μM) and chloroquine (EC_50_=1.13 μM) ^22, 23^. We have, therefore, further explored the antiviral efficacy of aglycone polyether ionophores against a broad-spectrum of viruses.

**Fig. 6.**
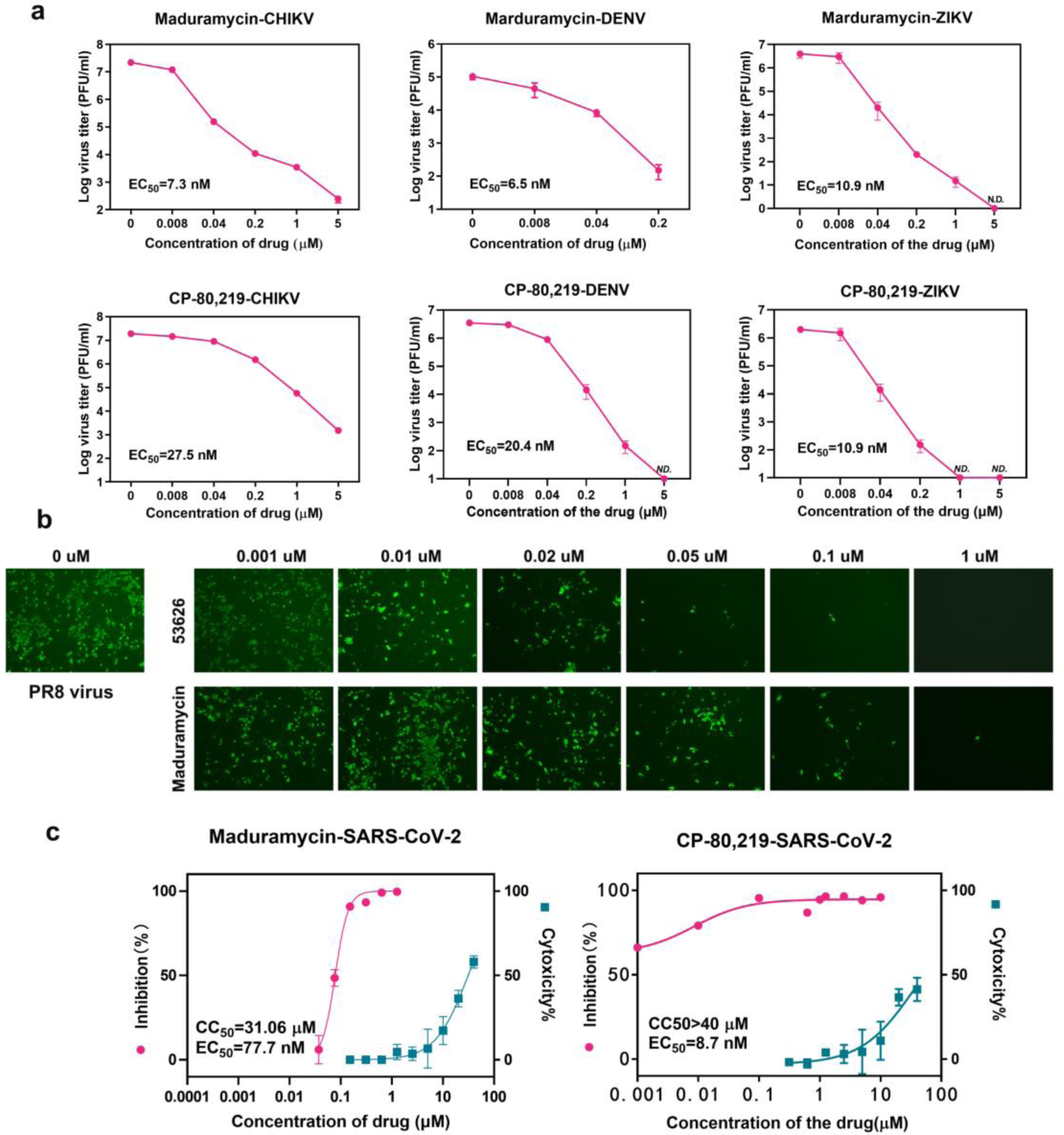
Inhibitory activity of maduramycin and CP-80,219 on different viruses. a) Dose-response analysis of maduramycin and CP-80,219 for CHIKV (n=5), DENV (n=5) and ZIKV (n=5). Vero cells were infected viruses with different concentrations of maduramycin or CP-80,219. The antiviral efficiency was evaluated by plaque reduction assay. b) Dose-response analysis of maduramycin and CP-80,219 on H1N1 influenza virus PR8. MDCK cells were infected PR8 (MOI=0.1) with different concentrations of maduramycin or CP-80,219 for 24 h, then measured by Immunofluorescence assay used CR9114 against HA (Scale bar = 50 µm). c) Dose-response analysis of maduramycin and CP-80,219 on SARS-CoV-2. Vero E6 cells were infected with SARS-CoV-2 (MOI=0.05, n=3) in the treatment of different doses of the indicated Maduramycin or CP-80,219 for 24 h. The viral yield in the cell supernatant was then quantified by qRT-PCR. Cytotoxicity of these drugs to Vero E6 cells was measured by CCK-8 assays (One representative data was shown from three independent experiments.). And all results are mean of three independent experiments.

## Discussion

Polyether ionophores, a very important class of natural products produced exclusively by actinomycetes, have the ability to selectively chelate metal ions (Na^+^, K^+^, Ca^2+^) and transport them across cell membranes to control coccidiosis^24^. These molecules, including maduramycin, monensin and salinomycin, are used as effective veterinary medicines and food additives in animal husbandry^25, 26, 27^. Recently, polyether ionophores have been shown to exert a broad-spectrum of bioactivity, including antiviral properties^28, 29, 30, 31, 32^. Here, we found that maduramycin and CP-80,219 aglycone polyether ionophores exhibit effective broad-spectrum antiviral activity for enveloped viruses, including JEV, ZIKV, DENV, CHIKV, and were also shown to exert promising active against influenza virus PR8 and SARS-CoV-2 pandemic coronavirus.

Many enveloped viruses enter cells via membrane fusion driven by low pH-induced conformational changes of specific viral proteins^33, 34, 35^. Studies have shown that monensin can affect glycosylation and transport of HIV envelope glycoproteins^17^ and alter vRNA trafficking^36^. In addition, monensin inhibits HCV entry by acting on endosome acidification^18^, similar to the mechanism of salinomycin against influenza virus^19^. Meanwhile, in the current study, we found that maduramycin did not decrease the pH value of cells (Supplementary Fig. 1), this indicated that aglycone polyether ionophore may not inhibit the fusion between enveloped viruses and cellular membranes by inducing alkalization of cells. Subsequently, cells are incubated maduramycin and JEV for 2 h, no difference was observed in the viral load of treated or untreated cells, further indicating that virus attachment and cell entry were not inhibited by maduramycin, a finding that was not in accordance with the results obtained for nanchangmycin, which was shown to impair cell entry of ZIKV^21^.

However, the virus titer in the supernatant was significantly reduced following addition of maduramycin after virus infection, suggesting that it might affect replication, assembly, or release of JEV. To confirm our hypothesis, we examined the viral nucleic acid load in cells treated with maduramycin and found that it was remarkably reduced after 12 h compared with the untreated group (Fig. 5d), Further, We found that maduramycin inhibited viral replication in the JEV replicon cell line in a time- and concentration-dependent manner (Fig. 5e), suggesting that aglycone polyether ionophores primarily interfere with JEV replication cycle. However, no inhibition was observed in the assembly and release model after exposure to maduramycin (Supplementary Fig. 3). Taken together, these results indicate that aglycone polyether ionophores elicit antivral effects against inhibit enveloped viruses via inhibition of a critical replication step in the replication cycle.

Moreover, considering that oral administration of salinomycin reportedly has no inhibitory effect on influenza virus, but rather induces potential toxicity and side effects^19^. We nest sought to further investigate the antiviral effects of maduramycin and CP-80,219 *in vivo*. To this end, we prepared liposome formulations based on the liposolubility of maduramycin and CP-80,219. Accordingly, we observed that their liposomes formulations were effective against JEV *in vitro*, and when administered *in vivo*, both maduramycin and CP-80,219 liposomes provided full protection of mice from JEV infection. In fact, virus titers in the blood were significantly decreased after 24 h of the first treatment (Fig. 4c). These data demonstrate that aglycone polyether ionophores encapsulated in liposomes have promising antiviral activity in mammals and may provide an effective strategy for other viruses that are currently epidemic in poultry, such as avian influenza viruses H1N1 and H7N9, as well as JEV and swine flu virus in pigs Drugs that are currently in use have great potential for being developed into antiviral drugs used in poultry and animal husbandry, which is important to avoid becoming a possible transmission or storage host and block the transmission between humans and animals.

In summary, we successfully demonstrated the antiviral activity of aglycone polyether ionophores in JEV-infected mice. Although, the diversity of viruses, and their associated pathogenic mechanisms, can affect the development of antiviral drugs, aglycone polyether ionophores are capable of exhibiting broad-spectrum antiviral activity, and probably target the functions of host cells to inhibit virus replication. However, further research is needed, particularly in elucidating the underlying mechanism of action, to facilitate the development of polyether ionophores into efficient antiviral drugs. Importantly, polyether ionophores can readily achieve very high yields to meet an increased demand, and their preparation has a low associated cost. For example, the yield of maduramicin titer can reach 7.16 g/L in shake-flask fermentation^37^. In short, the application of maduramycin and CP-80,219 aglycone polyether ionophores against JEV provides new insights for the development of antiviral drugs used in the poultry and animal husbandry industry, and brings hope for the development of broad-spectrum antiviral drugs.

## Materials and methods

### Cells, virus, compounds and antibody

BHK-21 (baby hamster kidney cell line), Vero (African green monkey epithelial kidney cell line), MDCK (Mardin–Darby canine kidney) and Vero E6 cells (ATCC, no.1586) were cultured in Dulbecco’s modified Eagle’s medium (DMEM; Invitrogen) with 10 % FBS, 100 U/mL of penicillin and 100 µg/mL of streptomycin at 37 °C with 5 % CO_2_.

JEV virus was derived from the infectious cDNA clone of pACYC-JEV-SA14^38^. The Asian lineage ZIKV isolate SZ-WIV001 (KU963796) was isolated from a Chinese patient in 2016^39^. The Asian genotype CHIKV (GenBank accession No. KC488650) was isolated from a clinically CHIKV-positive patient in China through seven rounds of serial passages in C6/36 cells^40^, PR8 (A/Puerto Rico/8/34) were stored at −80 °C in our laboratory and propagated in chicken embryos for use in all experiments. SARS-CoV-2 (IVCAS 6.7512) was propagated on the Vero-E6 cells and titrated by single layer plaque assay with standard procedure. Antibody-CR9114 against the HA protein of influenza A or B virus was kindly provided by Dr. Chen, jianjun.

### Preparation of 39 supernatant samples and six polyether ionophores

13 kinds of industrial strains of actinomycetes were provided by J1 Biotech Co. Ltd. (Wuhan, Hubei, China) and cultured on soybean flour-mannitol (SFM) agar plates [2 % (w/v) soybean flour, 2 % (w/v) mannitol, and 2 % (w/v) agar], The mycelium seeds of actinomycetes were grown in 50 mL of trypticase soy broth (TSB) for 3 days at 30 °C and agitated at 220 rpm, then pipetted 10 mL seed culture into 50 mL of the A fermentation medium [0.4 % (w/v) corn syrup, 6 % (w/v) glucose, 2.4 % (w/v) feather powder, 0.005 % (w/v) Fe_2_ (SO_4_)_3_, 0.3 % (w/v) NaCl, 0.015 % (w/v) K_2_HPO_4_, 0.1 % (w/v) CaCO_3_, pH 7.2], or B fermentation medium (soluble starch 30 g/L, soybean powder 10 g/L, yeast extract 2.5 g/L, calcium carbonate 3 g/L, and pH 7.2), or the C fermentation medium(3 % glucose, 1 % casein hydrolysate, 0.2 % sodium chloride, 0.2 % potassium chloride, 0.5 % ammonium sulfate, 0.02 % dipotassium hydrogen phosphate, 0.01 % magnesium sulfate, 0.01 % calcium chloride, 0.5 % calcium carbonate) for 7 days at 30 °C and 220 rpm.

The preparation of four polyethers (A-130-A, CP-80,219, maduramycin and edusamycin) were referred in our previous studies^9, 37, 41^. These compounds were purified using preparative reversed-phase high-performance liquid chromatography, maduramycin was prepared by recrystallization. Nanchangmycin and salinomycin have been prepared and reported by our lab^42, 43^. All drugs were dissolved in DMSO (Sigma-Aldrich) to make a 10 mM stock solution.

### Aglycone polyethers liposome preparation

The liposome was charged with a bilayer composed of Soybean and cholesterol at 4:1 weight ratio, then mixed in the organic solvent and evaporated using a dry nitrogen or rotary evaporation, and dissolve with aqueous dispersion (25 mL contains 2125 mg sucrose, 94 mg glycine and 7 mg calcium chloride dihydrate). The crude liposome disrupts using sonic energy (sonication), and then liposome extrusion is forced through a polycarbonate filter with a defined pore size to yield particles, average size of particles is measured by Malvern DSL.

### Antiviral activity assays of polyethers or supernatants

RT-PCR Assay. The cytotoxicity of the tested drugs on Vero E6 was determined by CCK8 assays (Beyotime, China). To evaluate the antiviral efficacy of these drugs, Vero E6 cells were cultured overnight in 48-well cell-culture petri dish with a density of 5 × 10^4^ cells/well. Cells were pre-treated with the different doses of the indicated antivirals for 1 h, and the virus (MOI of 0.05) was subsequently added to allow infection for 2 h. Then, the virus-drug mixture was removed and cells were further cultured with fresh drug-containing medium. At 48 hpi, the cell supernatant was collected and lysed in lysis buffer (Takara, Cat no. 9766) for further quantification analysis. One hundred microliter cell culture supernatant was harvested for viral RNA extraction using the MiniBEST Viral RNA/DNA Extraction Kit (Takara, Cat no. 9766) according to the manufacturer’s instructions. RNA was eluted in 30 μL RNase-free water. Reverse transcription was performed with a PrimeScript RT Reagent Kit with gDNA Eraser (Takara, Cat no. RR047A) and qRT-PCR was performed on StepOne Plus Real-time PCR system (Applied Biosystem) with TB Green Premix Ex Taq II (Takara Cat no. RR820A). Briefly, 3 μL total RNA was first digested with gDNA eraser to remove contaminated DNA and then the first-strand cDNA was synthesized in 20 μL reaction and 2 μL cDNA was used as template for quantitative PCR.

### Cytotoxicity assay

Cells were seeded in 96-well plates (1 × 10^4^ cells per well) and allowed to grow for 16 h before treatments. Afterward, compounds at concentrations in a 2-fold dilution series were added to the cell. At 36 h, the cells were incubated with 10 μL CCK8 reagent (cell counting kit-8, Bimake) for 1 h at 37 °C. The absorbance at 450 nm was read by a Microplate Reader (Varioskan Flash, Thermo Fisher). Cell viability was expressed as a percentage of the treated cells to the control (untreated) cells. For each compound concentration, six wells were performed in parallel, mean values of the cell viability were calculated. The CC50 was calculated by nonlinear regression using GraphPad Prism 8.0 software to determine the cytotoxic concentration at which 50 % of the cells are viable.

### Single-layer plaque assay

BHK-21 cells were seeded into 24-well plates at a density of 1 × 10^5^ cells per well one day before plaque assay. A series of 1:10 dilutions were made by mixing 15 µL of virus sample with 135 µL of DMEM. Then, 100 µL of each dilution were added to individual wells of 24-well plates containing confluent BHK-21 cells. The plates were incubated at 37 °C with 5 % CO_2_ for 1 h before the layer of 2 % methyl cellulose was added. After 3 days of incubation at 37 °C with 5 % CO_2_, the cells were fixed with 3.7 % formaldehyde and then stained with 1 % crystal violet. Plaque morphology and numbers were recorded after washing the plates with tap water.

### Times of addition assay

In this experiment, a monolayer of Vero cells was grown in 12-well plate in DMEM containing 2% inactivated FBS. The wells were designated as −1, 0,1, 2, 4, 8,10-h, that they are named according to the time of JEV-WT (MOI = 10) infection. As for −1 h, Vero cells were treated with marduramycin (500 nM). The plate was then incubated for 1 h at 37 °C with 5 % CO_2_. At the 0 h, all wells except the control wells were infected with JEV and again incubated for 1 h at 37 °C with 5 % CO_2_. After 1 h, the infected cells were washed thrice with PBS, and then 1 mL fresh medium was added to each well, and marduramycin (500 nM) were added to 1 h, 2 h, 4 h and 8 h wells respectively. The plate was then incubated at 37 °C with 5 % CO_2_ for 12 h. The viral titer was detected as described above plaque assay.

### Western blot analysis

Equal amounts of WT and mutant NS4A–NS4B-FLAG expression plasmids were mock-transfected or co-transfected with NS2B-3 pro-expression plasmid into 293T cells using Lipofectamine 2000 as described above. At 12 h p.t, cells were lysed with 200 mL lysis buffer containing 20 mM Tris (pH 7.4), 100 mM NaCl, 0.5 % n-dodecyl b-D-maltoside (Sigma) and EDTA-free protease inhibitor cocktail (Roche). Cell lysates were centrifuged at 17000 g for 20 min, and the supernatants were collected and heated at 75 °C for 10 min. Samples were separated using 15 % SDS-PAGE gels and transferred to PVDF membranes (Millipore), followed by blocking with 5 % skim milk (Bio-Rad) in TBST buffer (50 mM Tris/HCl, pH 7.5, 150 mM NaCl and 0.1 % Tween 20) at room temperature for 1 h. Following blocking, membranes were subject to sequential incubation with anti-FLAG mouse antibody (1: 2000 dilution) and secondary anti-mouse IgG conjugated to HRP (Bio-Rad). After washing three times with TBST buffer, the signals were detected with a chemiluminescence system (ChemiDoc; Bio-Rad).

### In vivo experiments

We performed mouse studies to evaluate the *in vivo* efficacy of maduramycin and CP-80,219. Female sex C57BL/6 mice, 3–4 weeks old, were kept in biosafety level 2 laboratory and given access to standard pellet feed and water ad libitum. All experimental protocols followed the standard operating procedures of the approved biosafety level 2 animal facilities. JEV infection in C57BL/6 mice (n = 5/dose group) by subcutaneously inoculated with 1 × 10^7^ PFU in 200 μL PBS. After infection 30 min, Vehicle or liposomes (1 mg/kg) are administered intraperitoneally daily, and the weight was measured daily from 0 to 14 days post infection (dpi). Survival and general conditions were monitored for 14 days or until death. Blood was collected daily from 1 to 3 dpi to track for viral titration via plaque assay. All animal experiments were performed according to protocols and ethical guidelines approved by Center for Animal Experiment and BSL-3 laboratory at Wuhan Institute of Virology.

### Real time RT-PCR

To detect the copy number of RNA replication in cells, total cellular RNA was isolated using the Qiagen RNeasy Mini Kit (QIAGEN), and subjected to SYBR green one-step real-time RT-PCR as described previously44. The copy number of replicons was calculated by interpolating in a standard curve made by a quantified JEV replicon plasmid44.

For inhibitory assay of SARS-CoV-2, RNA was extracted from 140 μL cell supernatant using QIAamp viral RNA mini kit (52906, Qiagen) following the manufacturer protocol. QRT-PCR were performed using Luna® Universal Probe One-Step RT-PCR Kit (E3006). Average values from duplicates of each gene were used to calculate the viral genomic copies. RBD-qF1: 5’-CAATGGTTTAACAGGCACAGG-3’; RBD-qR1: 5’-CTCAAGTGTCTGTGGATCACG-3’; Probe: ACAGCATCAGTAGTGTCAGCAATGTCTC.

### Western blot analysis

Cells were lysed with 200 µL of lysis buffer containing 1 % Triton X-100, 50 mM Tris–HCl (pH=8.0), 150 mM NaCl, 1 mM EDTA and 1 mM PMSF on ice for 10 min. Following boiling at 95 °C for 10 min, cell lysates were separated by 10 % SDS-PAGE. Separated proteins were then electro-transferred onto a polyvinylidene fluoride (PVDF) membrane (0.2 µm; Millipore). After blocked with 5 % skim milk in TBST (50 mM Tris-HCl, 150 mM NaCl, 0.1 % Tween 20, Ph=7.4) for 1 h at room temperature, the membranes were then incubated with primary antibodies at room temperature for another 1 h. After washing three times with TBST, the membranes were incubated with horseradish peroxidase (HRP) conjugated goat anti-rabbit or mouse secondary antibody (Bio-Rad) for 1 h at room temperature, followed by washing three times with TBST. The protein bands were visualized with a chemiluminescent detection reagent (Millipore).

### Statistical analysis

All statistical data analyses were performed in Graphpad Prism 8. Statistical significance for each endpoint was determined with specific statistical tests. In general, for metrics with multiple treatment groups with longitudinal data (e.g., mouse weight loss or pulmonary function over time), two-way ANOVA was performed with the suggested multiple comparison test as advised by Prism. For comparative data with for a single timepoint (e.g., lung titer) Kruskal-Wallis or one-way ANOVA was performed with the suggested multiple comparison test. For each test, a p-value <0.05 was considered significant. Specific tests are noted in each figure legend.

## Acknowledgements

The work was financially supported by the National Key R&D Program of China (2018YFA0900400), the National Natural Science Foundation of China (31670090), the Fundamental Research Funds for the Central Universities (2042020kf1003) and the Medical Science Advancement Program (Clinical Medicine) of Wuhan University (TFLC2018002). We thank J1 Biotech Co.,Ltd for providing all actinomycete strains, and also thank Wanyi Tai from Wuhan University School of Pharmaceutical Sciences for his help in the preparation of liposomes. The experiments related to SARS-CoV-2 were completed at National Biosafety Laboratory, Wuhan, Chinese Academy of Sciences. We are particularly grateful to Tao Du, Jin Xiong and Lun Wang from Zhengdian Biosafety Level 3 Laboratory and the running team of the laboratory for their work. We are grateful to the staffs from xiaohongshan (Xue-fang An, Fan Zhang, He Zhao and Li Li) and Zhengdian (YanFeng Yao, Jing Deng, Ge Gao and Yun Peng) Center for Animal Experiment at Wuhan Institute of Virology for their helpful supports during the course of the work.

## Author contributions

T.L. has the original idea. T.L. and B.Z. designed the experiments. M.H., L.R. and Y.W. provided all fermented samples and preparation of polyether ionophores. Q.J. performed and analyzed antiviral activity and cytotoxicity of all samples and polyether ionophores *in vitro* and *in vivo* against JEV, DENV, ZIKV and CHIKV. Y.N.Z. only performed and analyzed antiviral activity and cytotoxicity of maduramycin and CP-80,219 *in vitro* against SARS-CoV-2. Z.Z. and Q.Z performed antiviral activity of maduramycin and CP-80,219 *in vitro* against PR8 virus. M.H. B.Z. and T.L. analyzed the data. M.H., Q.J., B.Z., and T.L. wrote the paper.

