## supplementary information for "Aglycone polyether ionophores as broad-spectrum agents inhibit multiple enveloped viruses including SARS-CoV-2 in vitro and successfully cure JEV infected mice"

| Industrial actinomycete strains | Main products | Types of antibiotic |
|---|---|---|
| J1-002 | Natamycin | macrolide |
| J1-005 | Ɛ-poly-lysine | polylysine |
| J1-007 | Maduramycin | polyether |
| J1-008 | Salinomycin | polyether |
| J1-009 | Daptomycin | cyclic lipopeptide |
| J1-010 | Apramycin | aminoglycoside |
| M3 | Erythromycin | macrolide |
| J1-019 | Spinosad | macrolide |
| J1-021 | Tylosin | macrolide |
| J1-022 | Chlortetracycline | tetracycline |
| J1-B | Gentamicin B | aminoglycoside |
| J1-C | Gentamicin C | aminoglycoside |
| J1-A | Abamectin | macrolide |
| **Wild type actinomycete strains** | **Main products** | |
| J1-001 | Nanchangmycin | polyether |
| ATCC-21840 | A-130-A | polyether |
| ATCC-39574 | Endusamycin | polyether |
| ATCC-53626 | CP-80,219 | polyether |

**Supplementary Table 1. 13 industrial strains and 4 wild type actinomycete strains**

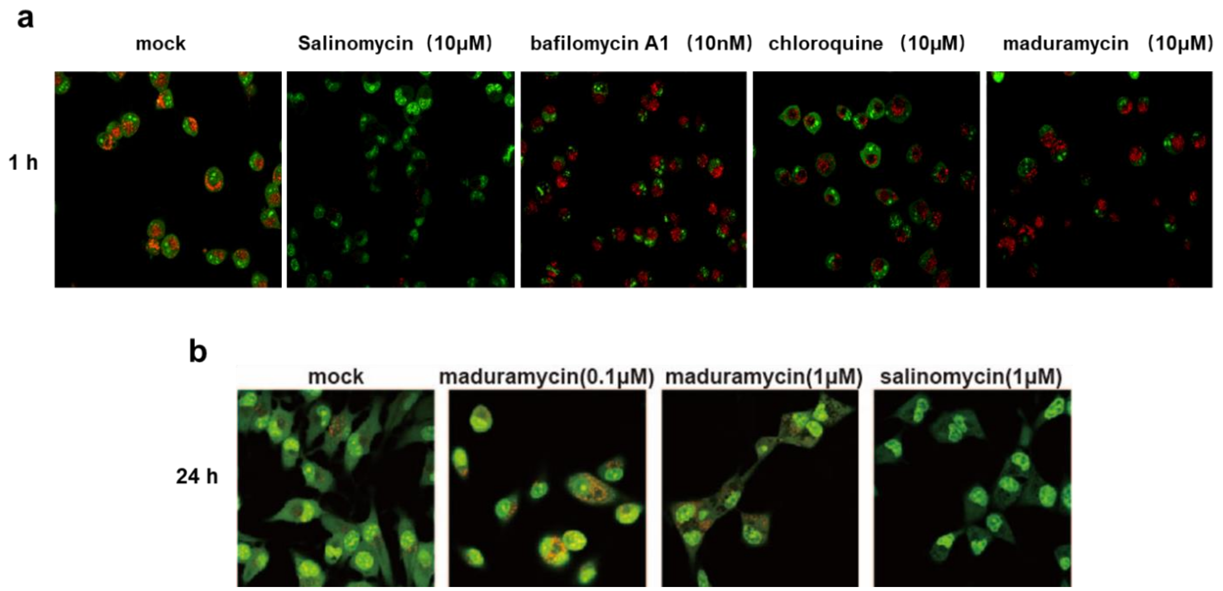


**Supplementary Fig. 1. Maduramycin did not affect endosome acidification to disrupt viral particles.** a) 10 μM maduramycin was premixed with PR8 virus for incubation at 37 °C for 1 h, salinomycin (10 μM) as positive control, bafilomycin (10 μM) and chloroquine (10 μM) as negative control. b) 1 μM maduramycin was premixed with PR8 virus for incubation at 37 °C for 24 h Vero cells then were treated with acridine orange (4 μg/ml) for 10 min and observed by confocal microscopy. Scar bar: 600 nm.

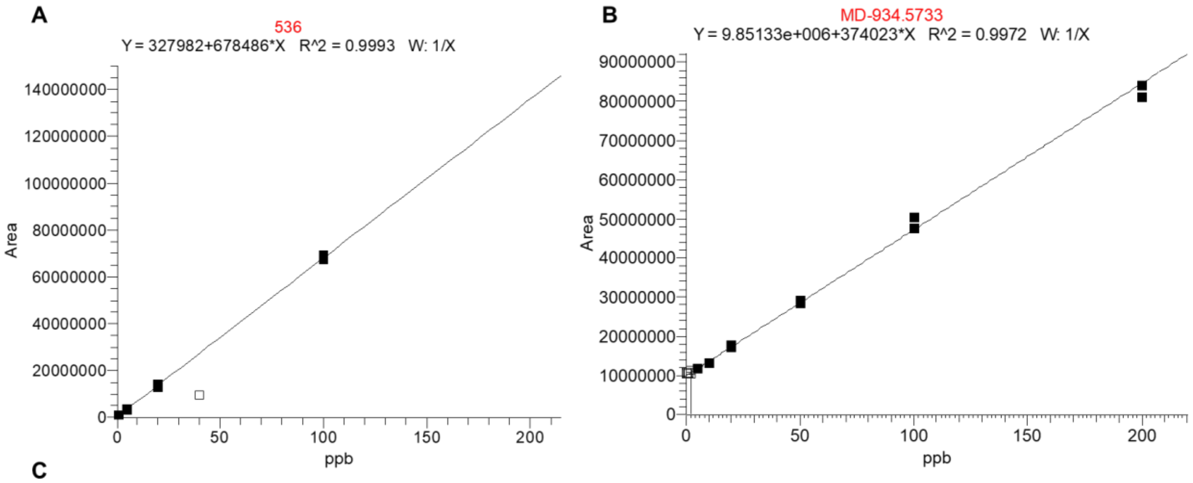


C

| Compounds | Liposome average size(nm) | Rate of encapsulation efficiency (%) |
|---|---|---|
| Maduramycin(MD) | 105 | 85.7 ±2.3 |
| CP-80,219(536) | 107 | 89.1 ±3.7 |

**Supplementary Fig. 2. Liposome encapsulation efficiency for maduramycin and CP-80,219.** A+B) The standard curves of maduramycin (MD) and CP-80,219 (536) were recorded by LCMS result, samples were dissolved in acetonitrile, loaded onto a C18 column (5 μm, 5 ×0.3 mm, Agilent Technologies, Inc.), and eluted at a flow rate of 0.2 mL/min to mass spectrometry (MS) analysis. C) the results of Liposome encapsulation efficiency for maduramycin and CP-80,219, Three replicates were used for each sample.

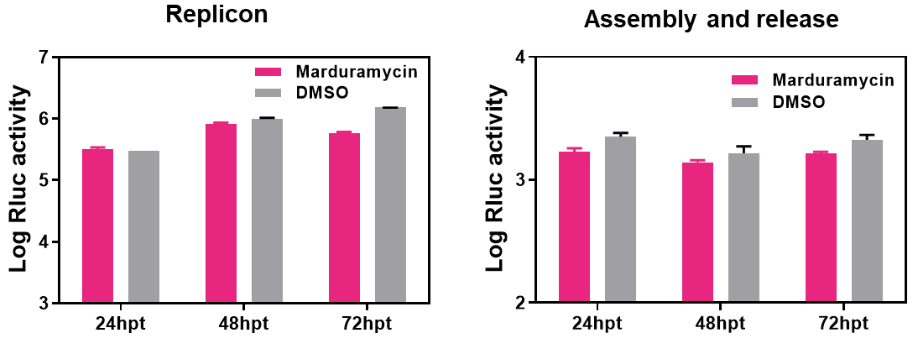


**Supplementary Fig. 3. Maduramycin not obviously effect JEV assembly and release.** We used JEV particles, which contain two RNA being responsible to JEV replication and structure proteins, to imitate JEV assembly and release process. Vero cells was infected JEV particles for 2 h, then added to maduramycin (500 nM) for 24 hpt (hour post transfection), 48 hpt and 72 hpt, then detected Rluc activity.

**Supplementary Fig. S4. High resolution mass spectrometry results of salinomycin**

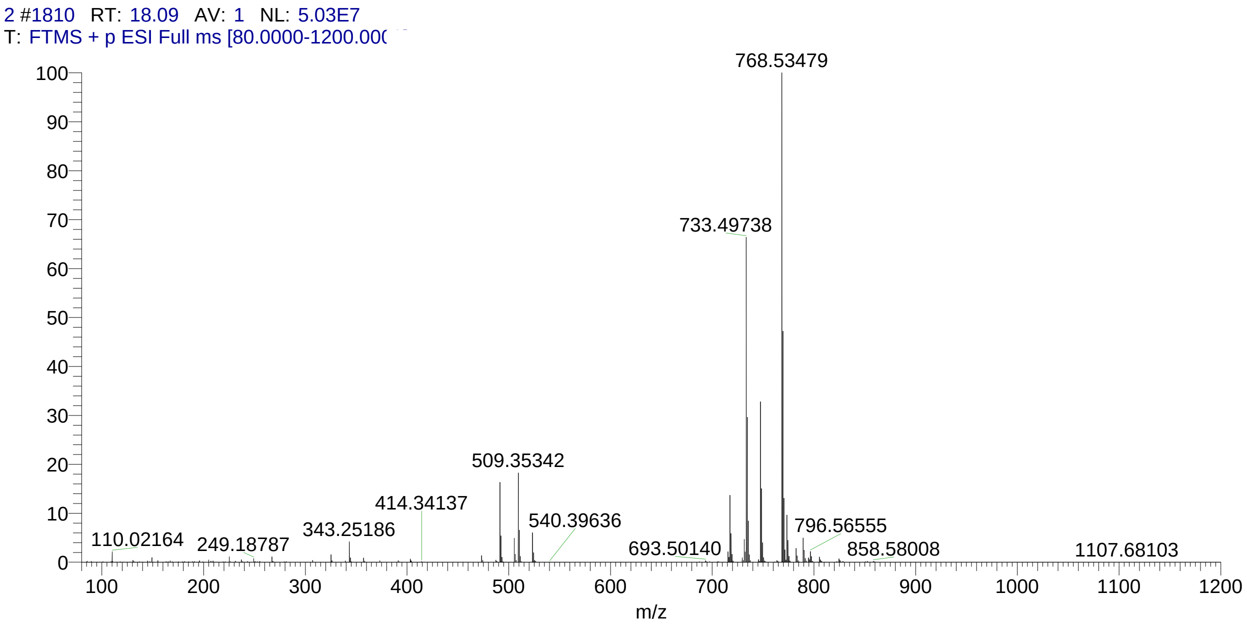


**Supplementary Fig. S5. High resolution mass spectrometry results of maduramycin**

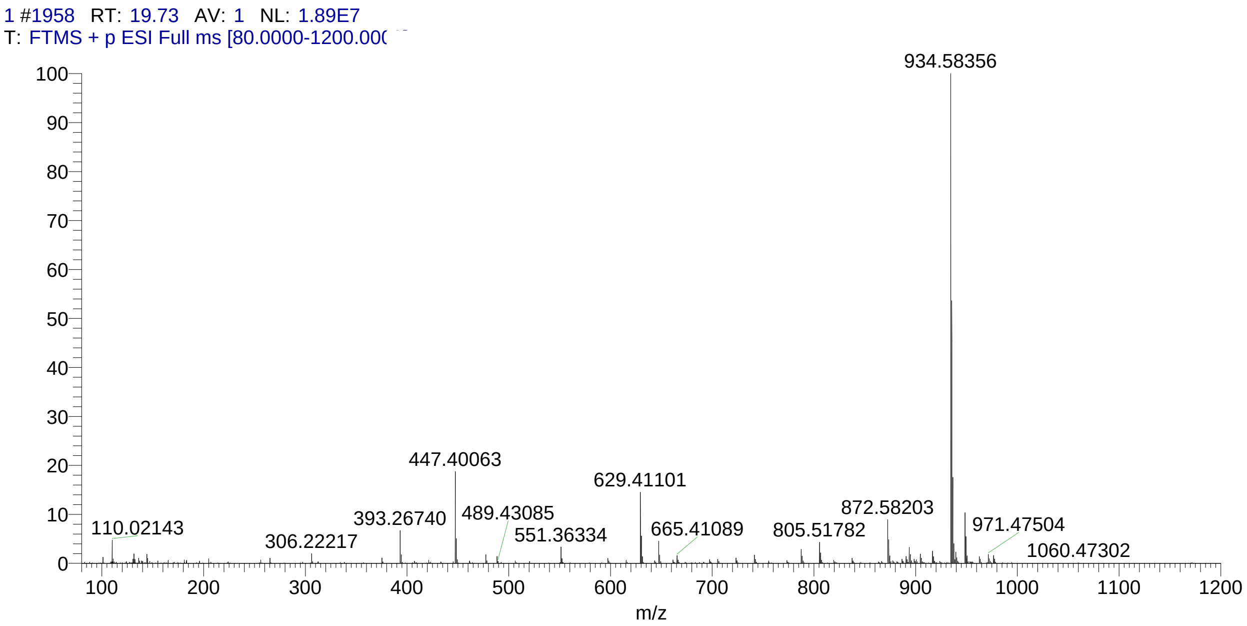

#### Supplementary Fig. S6. High resolution mass spectrometry results of CP-80,219

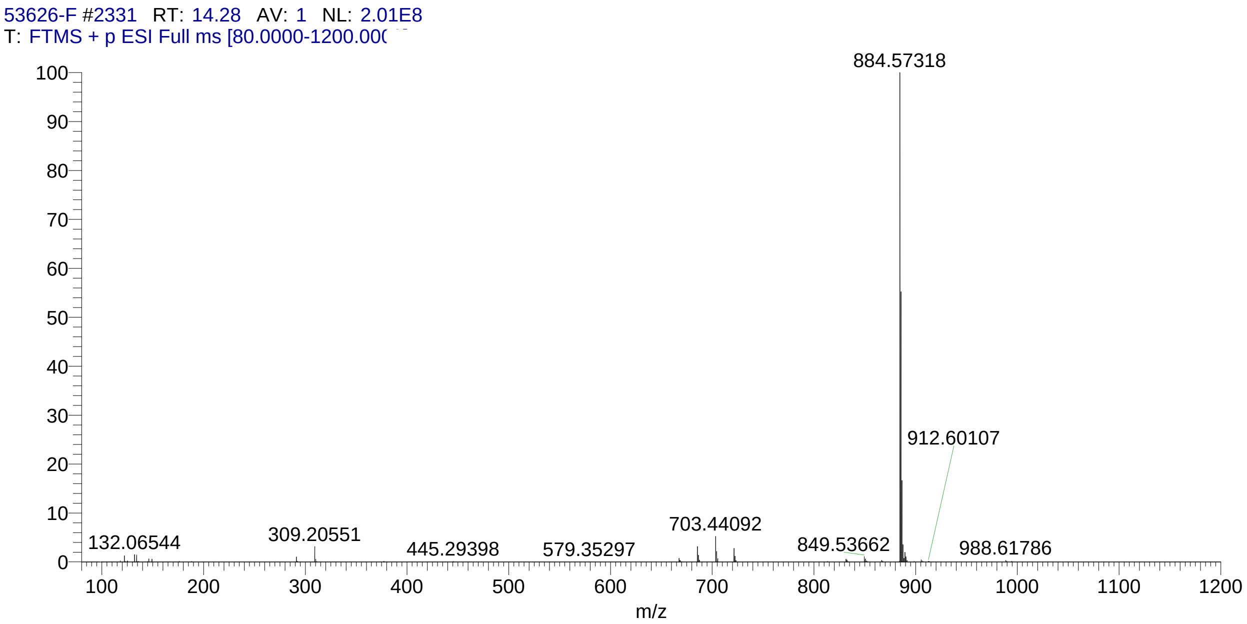


#### Supplementary Fig. S7. High resolution mass spectrometry results of endusamycin

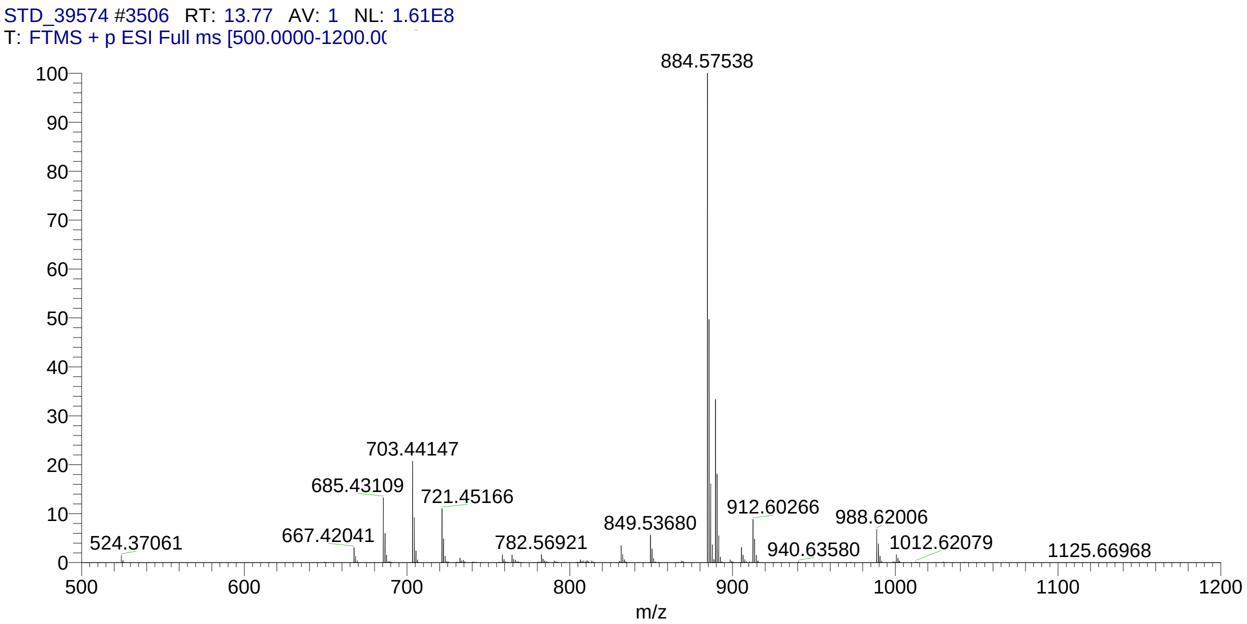

#### Supplementary Fig. S8. High resolution mass spectrometry results of nanchangmycin

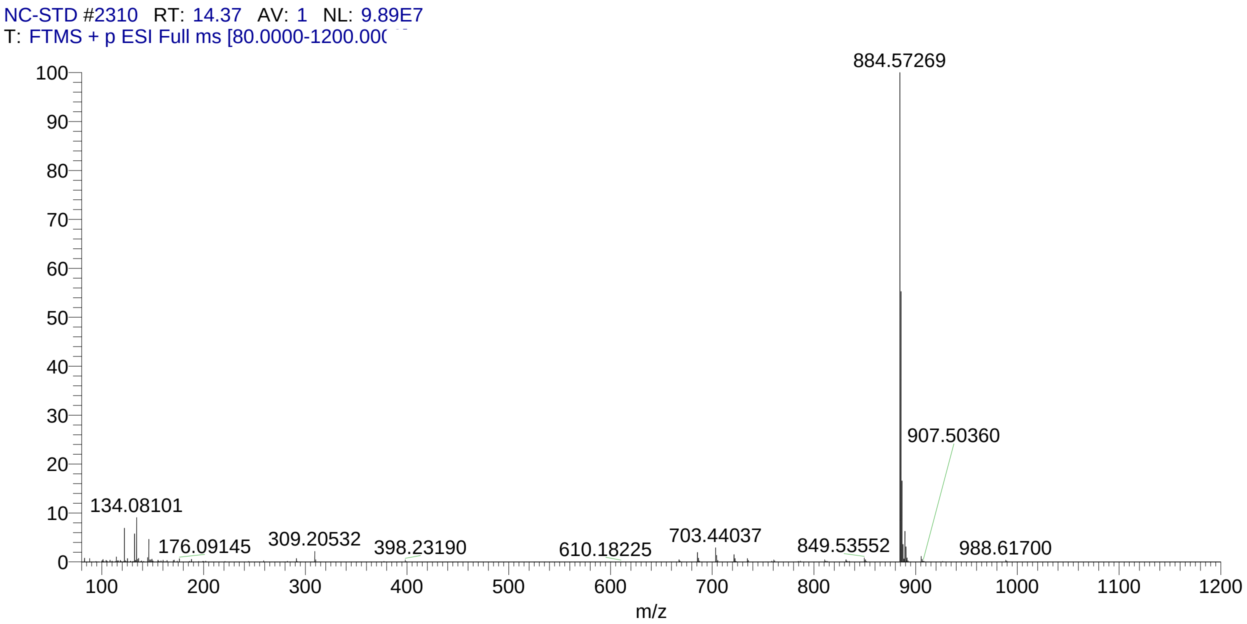


#### Supplementary Fig. S9. High resolution mass spectrometry results of A-130-A

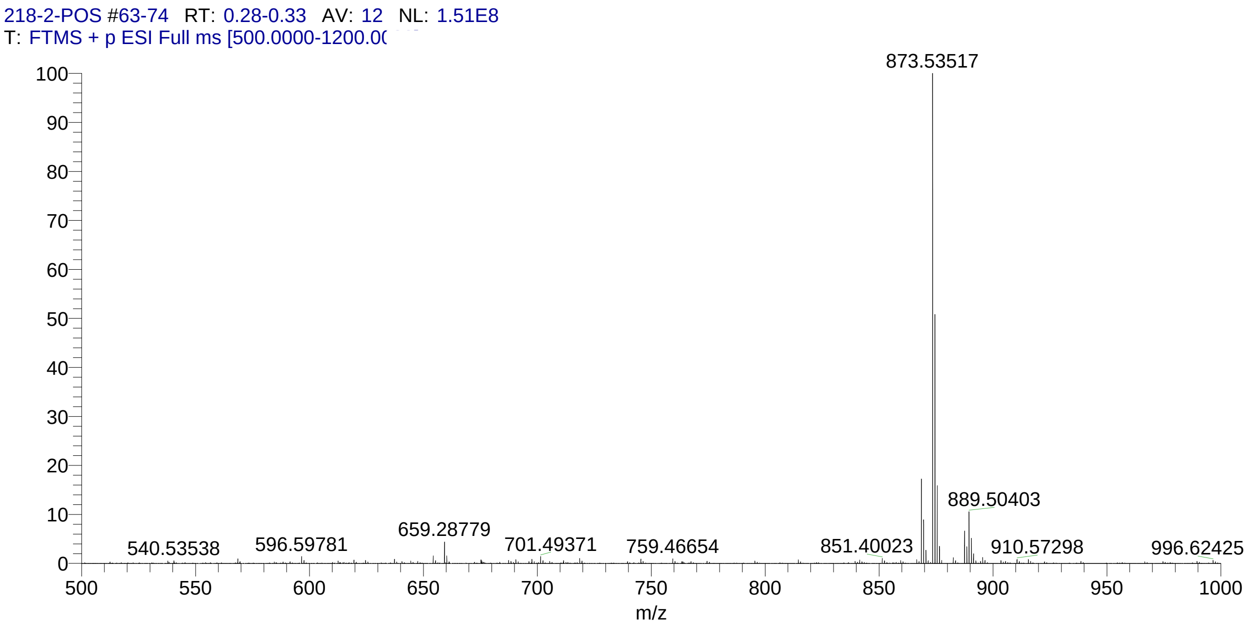

$^{1}H$ NMR of Maduramycin

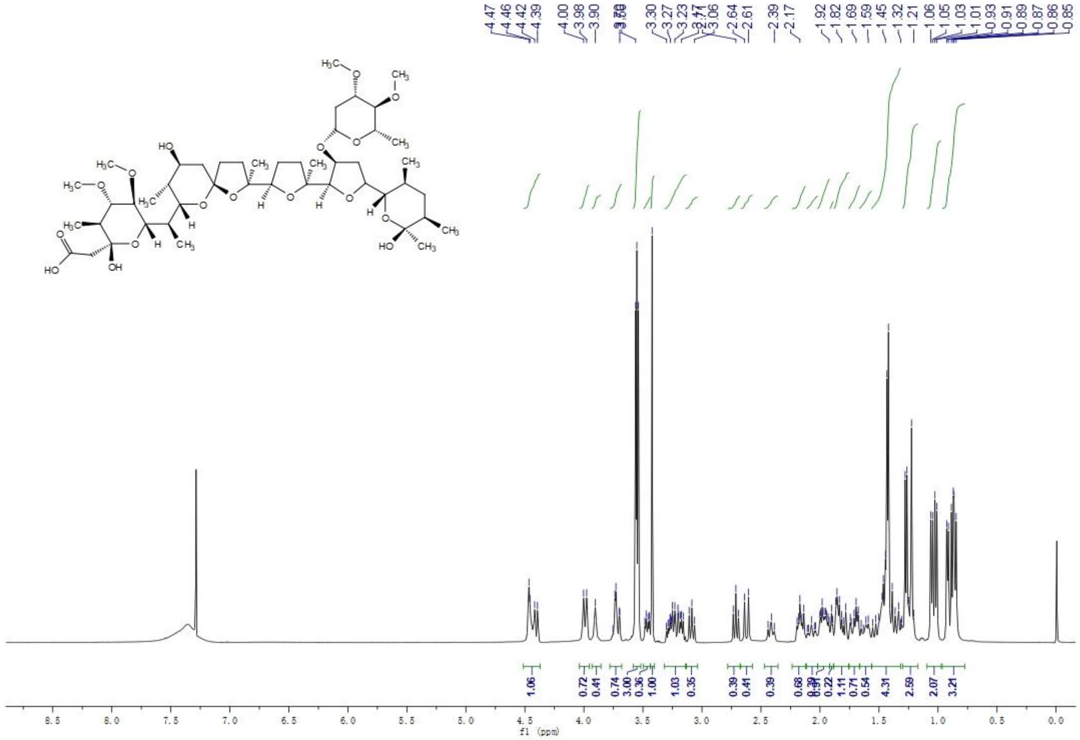

$^{13}C$ NMR of Maduramycin

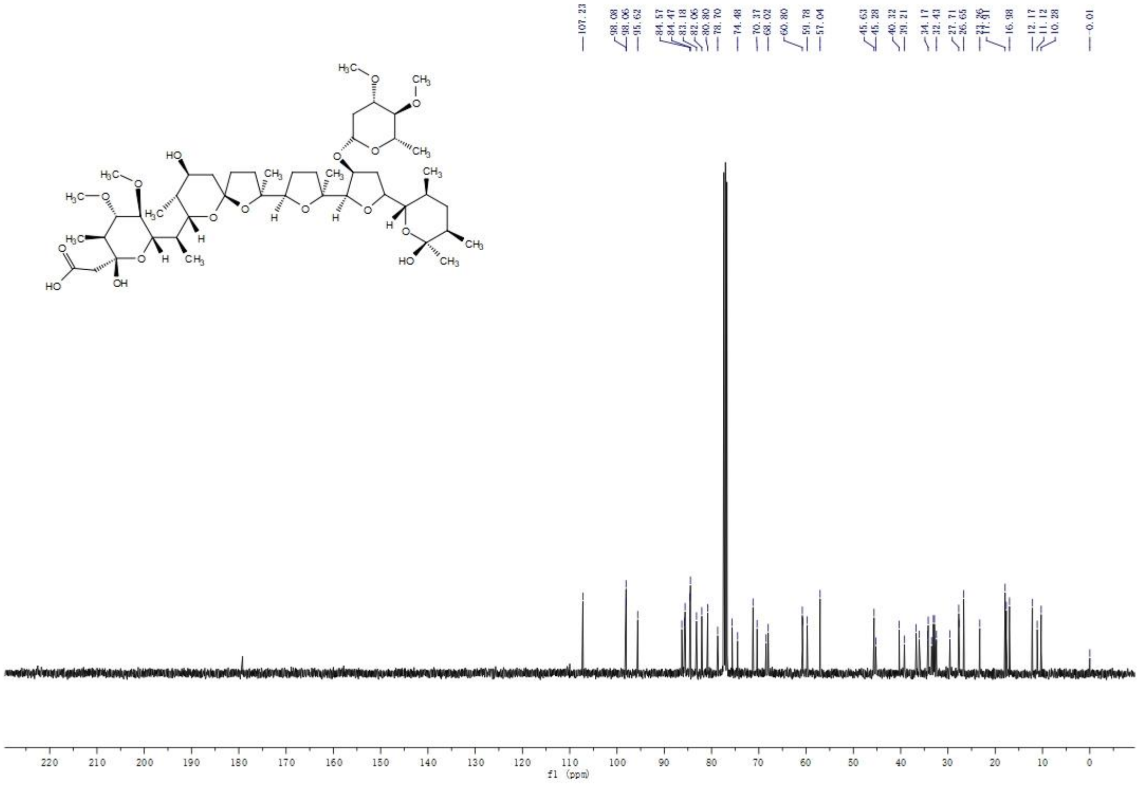

$^{1}H$ NMR of A-130-A

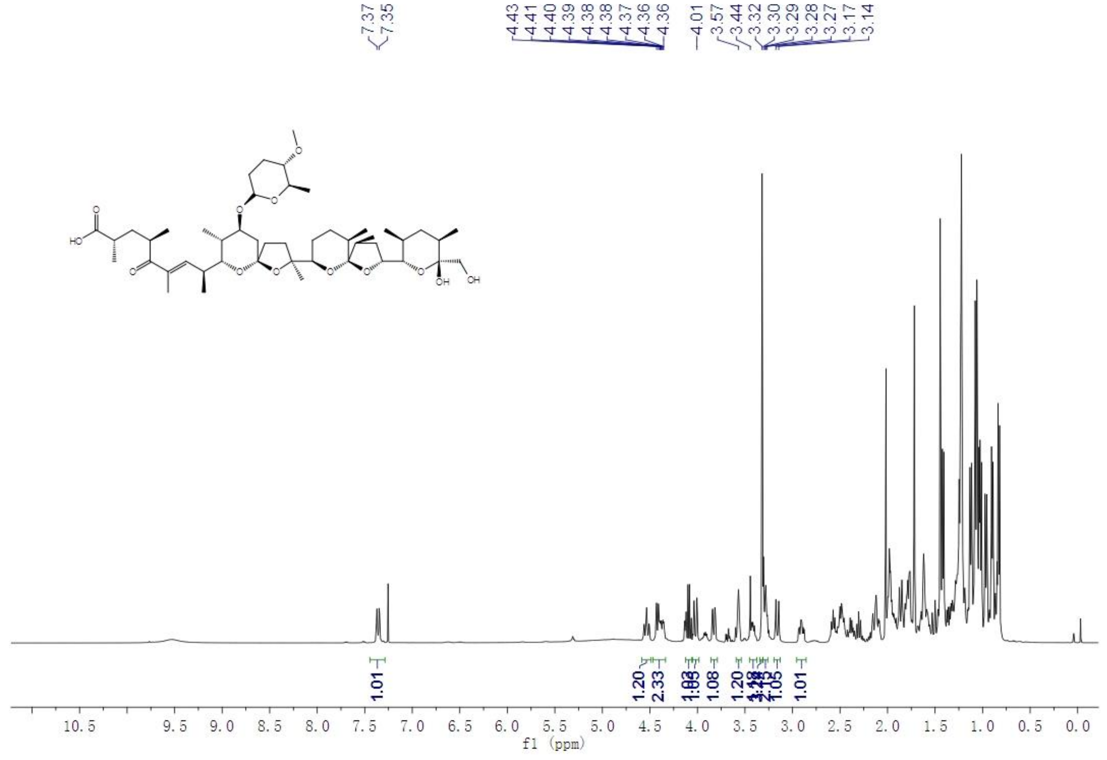

$^{13}C$ NMR of A-130-A

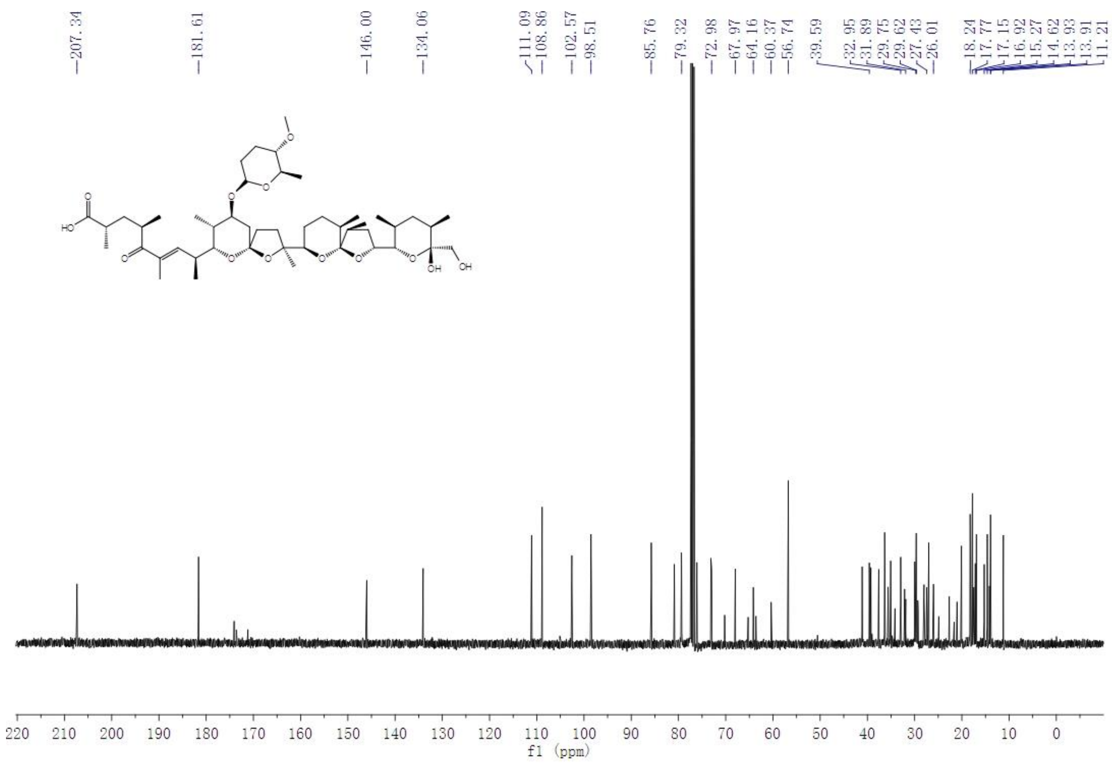

### $^{1}$H NMR of Endusamycin

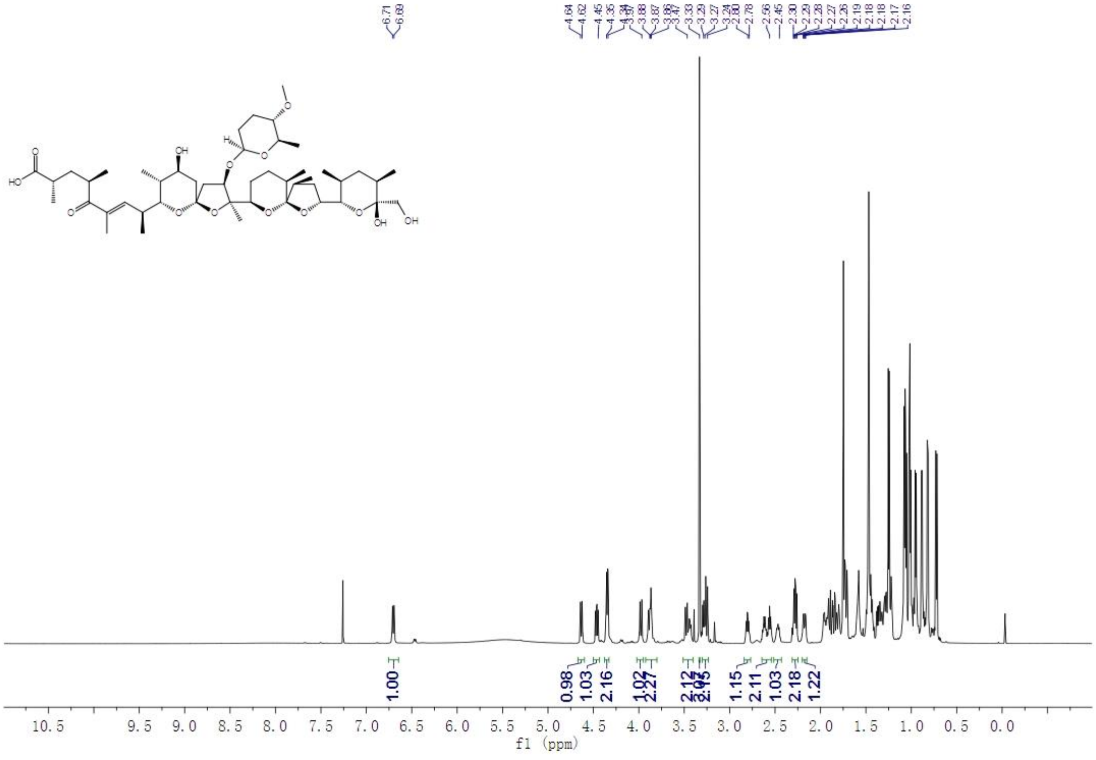


### $^{13}$C NMR of Endusamycin

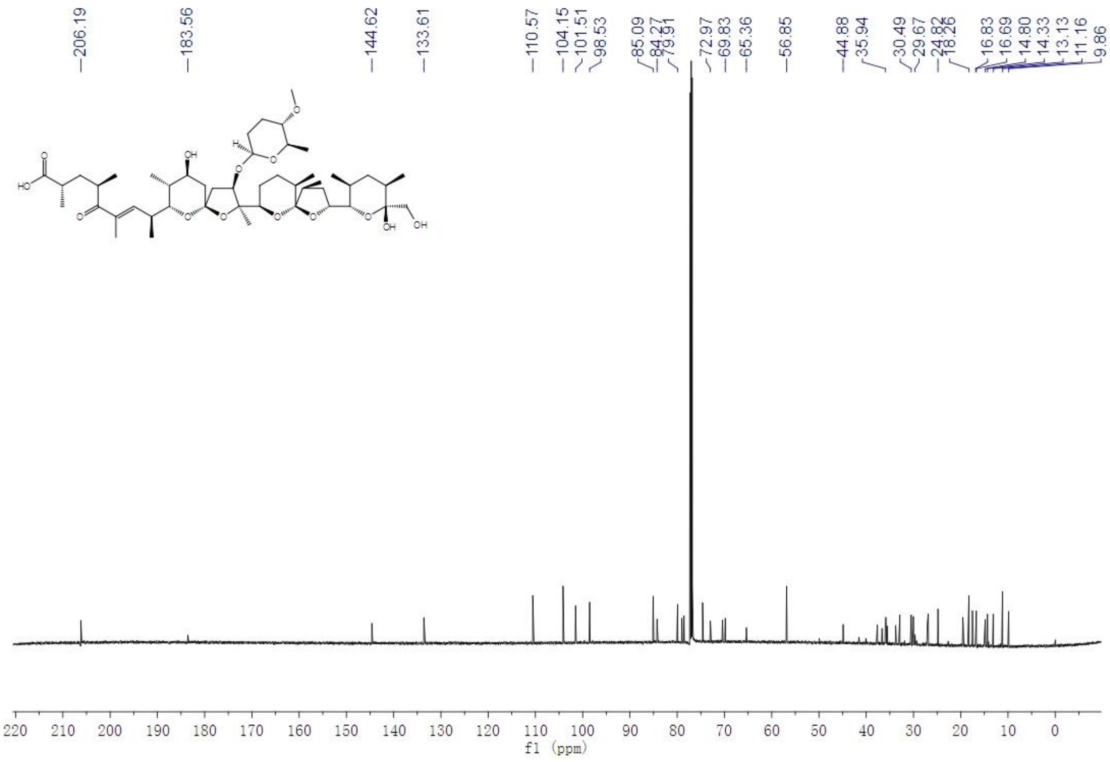

$^{1}H$ NMR of CP-80,219

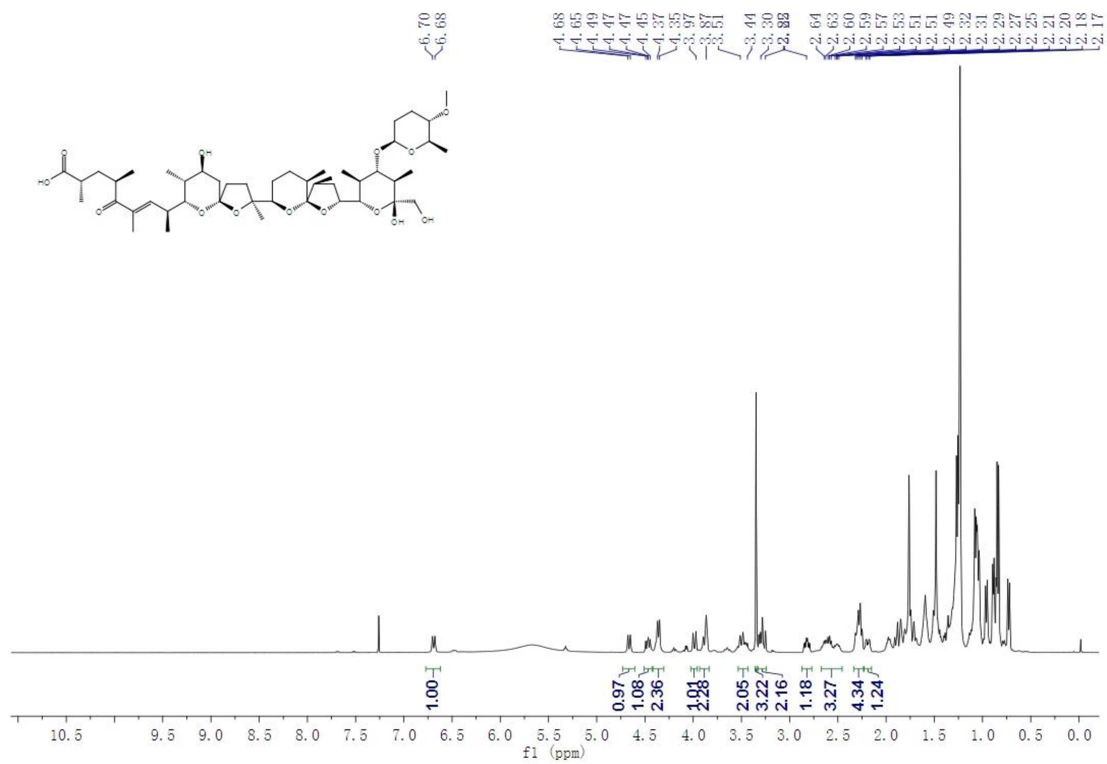

$^{13}C$ NMR of CP-80,219

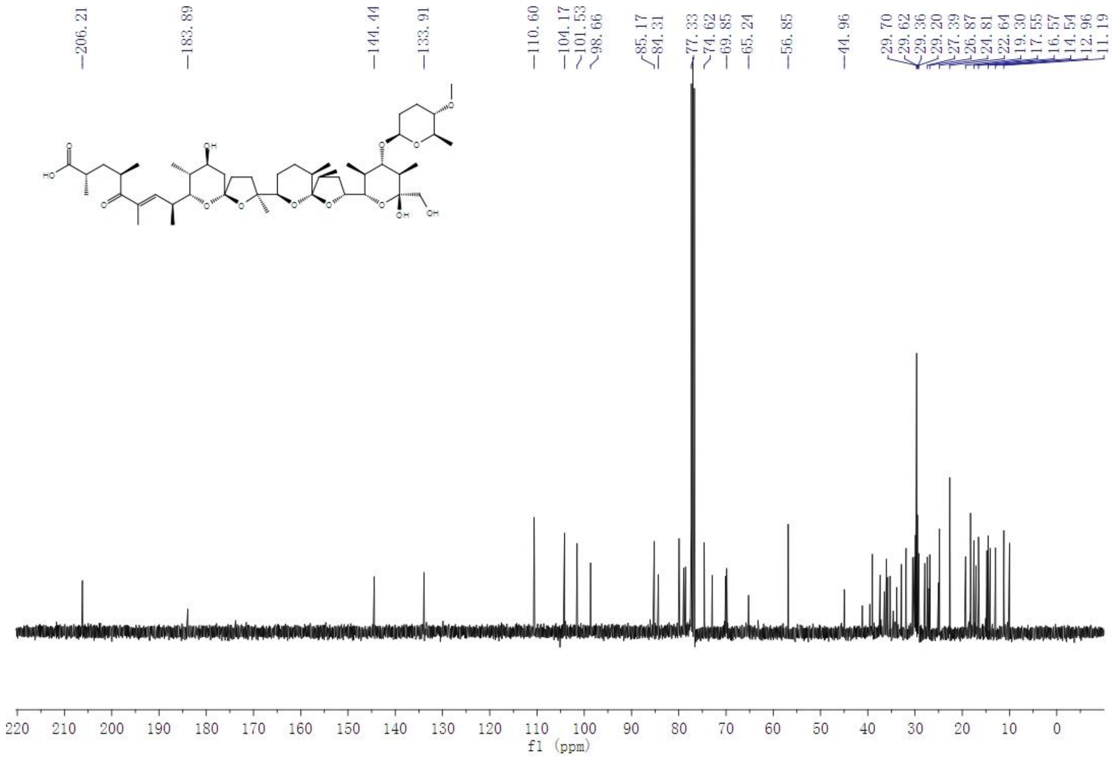
